# Association between chronotype and dual-task gait cost across distinct cognitive domains in healthy young adults

**DOI:** 10.64898/2026.04.16.719112

**Authors:** Meeyoung Kim, Jood Dalbah, Alham Jehad Ali Al-Sharman

## Abstract

Chronotype reflects individual circadian preference for timing of sleep, wakefulness, and peak performance and has been linked to variability in prefrontal cognitive function across the day. Whether chronotype independently relates to dual-task gait cost (DTC) and whether this relationship differs by cognitive task domain is unclear. Sixty-nine healthy young adults (37 female; mean age 21.3 years) completed the Morningness–Eveningness Questionnaire (MEQ). Spatiotemporal gait parameters were recorded with three-dimensional motion capture during single-task walking and three dual-task conditions: backward word spelling (5LWB; phonological), serial subtraction by seven (SS7; arithmetic), and reverse month recitation (RMR; sequential). DTC was calculated for eight gait parameters. Condition differences were assessed with nonparametric tests and post-hoc comparisons. Multiple linear regression, adjusting for age, sex, BMI, and baseline gait velocity, tested the independent association between MEQ score and mean velocity DTC; exploratory Spearman correlations examined other parameters. SS7 produced the largest mean velocity DTC (−12.76%), significantly greater than 5LWB (−7.95%; p = 0.002) and RMR (−9.57%; p = 0.021). MEQ score independently predicted mean velocity DTC in 5LWB (β = −0.51, p < 0.001, R² = 0.269) and RMR (β = −0.55, p = 0.004, R² = 0.222), indicating greater morningness associated with better gait-speed preservation under cognitive load; the SS7 association was not significant (β = −0.33, p = 0.071). Exploratory correlations showed MEQ–DTC associations across 7/8 parameters in 5LWB, 4/8 in RMR, and 3/8 in SS7. Chronotype is independently associated with dual-task gait cost in a task-domain-specific manner, with stronger effects for phonological and sequential tasks than for arithmetic processing. The SS7 condition yielded the largest interference but weakest chronotype modulation, suggesting arithmetic dual-task disruption may be less sensitive to circadian arousal. Fixed testing time and cross-sectional design warrant within-subject, multi-timepoint studies to confirm chronotype effects separate from time-of-day confounds.

## Introduction

Chronotype refers to the stable inter-individual variation in endogenous circadian phase, expressed as a preference for the timing of sleep, wakefulness, and peak physiological and cognitive performance [1,2]. It is shaped by genetic factors, including polymorphisms in core circadian clock genes, and is further modulated by age, sex, and socio-environmental zeitgebers such as light exposure and academic schedules [1,3]. The Morningness–Eveningness Questionnaire (MEQ) is a validated 19-item self-report instrument widely used to quantify this preference, classifying individuals along a continuum from definite evening to definite morning type [2,4]. In young adult populations, intermediate and evening chronotypes predominate, partly due to a biologically driven circadian phase delay in adolescence and early adulthood and partly due to social factors such as academic timetables and artificial light exposure at night [4,5].

A substantial body of evidence has established that chronotype influences cognitive performance in a time-dependent manner, consistent with the circadian synchrony hypothesis: performance is optimized when task timing aligns with an individual’s peak circadian arousal state [6,7]. Morning types typically reach this peak in the late morning to early afternoon, while evening types experience a delayed arousal trajectory with cognitive performance peaking later in the day [6,7]. The cognitive functions most sensitive to this circadian modulation include sustained attention, working memory, and executive control—processes underpinned by prefrontal neural networks—whereas more automatic or modality-specific processes are comparatively less affected [7,8].

Although walking is largely automatic in healthy young adults, it becomes attentionally demanding when performed concurrently with a secondary cognitive task [9,10]. The dual-task paradigm is widely used to quantify this cognitive–motor dependency: the change in gait spatiotemporal parameters relative to single-task walking is expressed as dual-task cost (DTC), an established index of cognitive–motor interference reflecting the extent to which finite attentional resources must be shared between locomotor control and concurrent cognitive processing [11,12]. Gait velocity, cadence, stride length, and stride and stance times are among the parameters most consistently sensitive to dual-task manipulation in healthy adults [11,12].

The magnitude and pattern of dual-task gait interference depend substantially on the cognitive domain engaged by the secondary task [13,14]. Tasks requiring phonological processing, working memory updating, or response inhibition—such as backward spelling or reverse sequential recitation—engage prefrontal executive networks that overlap substantially with the cortical circuits supporting gait under attentional load [15,16]. Tasks relying on arithmetic processing may additionally recruit distinct parietal circuits, potentially producing a different distribution of neural demand [15,16]. These differences in processing strategy, neural circuitry, and degree of overlap with locomotor control systems can yield distinct patterns of gait interference across task types [17,18]. Characterizing these domain-specific effects is therefore essential for interpreting individual variability in dual-task gait performance.

Despite the well-established influence of chronotype on prefrontal cognitive function and the known prefrontal contribution to dual-task gait regulation, the relationship between chronotype and dual-task gait performance has received limited empirical attention. Studies examining time-of-day effects on gait have documented diurnal variability in both single-task and dual-task walking performance [19,20], but few have assessed chronotype as a continuous individual-difference variable or systematically compared its effects across cognitive task domains with distinct neural demands. Given that chronotype modulates the availability of prefrontal arousal resources during a given testing window and given that different task domains engage these resources to varying degrees, chronotype effects on DTC would be expected to be task-domain-dependent rather than uniform [8,21]. Whether this is the case has not been directly examined.

Any investigation of chronotype effects on cognitive–motor performance must account for daytime sleepiness, which covaries with chronotype in young adult samples and could independently impair dual-task gait [3,5]. Evening types assessed during morning or early-afternoon windows may exhibit greater sleepiness relative to morning types, introducing a potential confound. Similarly, undetected variation in global cognitive status could influence dual-task gait independent of chronotype. The present study therefore incorporated the Epworth Sleepiness Scale (ESS) as an eligibility screen for pathological sleepiness and the Short Orientation-Memory-Concentration Test (SOMCT) as a cognitive screening instrument, minimizing the likelihood that observed associations reflect these factors.

This study aimed to examine the association between chronotype, measured continuously using the MEQ, and dual-task gait cost across three cognitive conditions representing distinct processing domains—phonological (backward word spelling; 5LWB), arithmetic (serial subtraction by seven; SS7), and sequential (reverse month recitation; RMR)—in healthy young adults. We hypothesized that: (1) dual-task gait cost would differ significantly across the three cognitive conditions, reflecting their differing neural demands; and (2) chronotype would be independently associated with dual-task gait cost in a task-domain-specific manner, with stronger associations in conditions engaging prefrontal executive and sequential processing networks (5LWB and RMR) than in the arithmetic SS7 condition.

## Materials and methods

### Study design and setting

This cross-sectional observational study investigated the association between chronotype and dual-task gait performance in healthy young adults. Data were collected at the Motion Analysis Laboratory, University of Sharjah, United Arab Emirates, between June 2023 and June 2024. All assessments were conducted within a fixed time window of 12:00–16:00. This window was adopted for logistical reasons and was not aligned to participants’ individual circadian preferences; its implications for interpreting the chronotype findings are discussed in the Limitations section.

### Participants

A total of 69 participants were recruited from a university population using convenience sampling (classroom advertisements). Eligibility screening was performed prior to enrolment. Participant flow is summarized in the study flowchart shown in **Fig 1**. (n = 70 assessed; n = 69 included).

**Fig 1.**
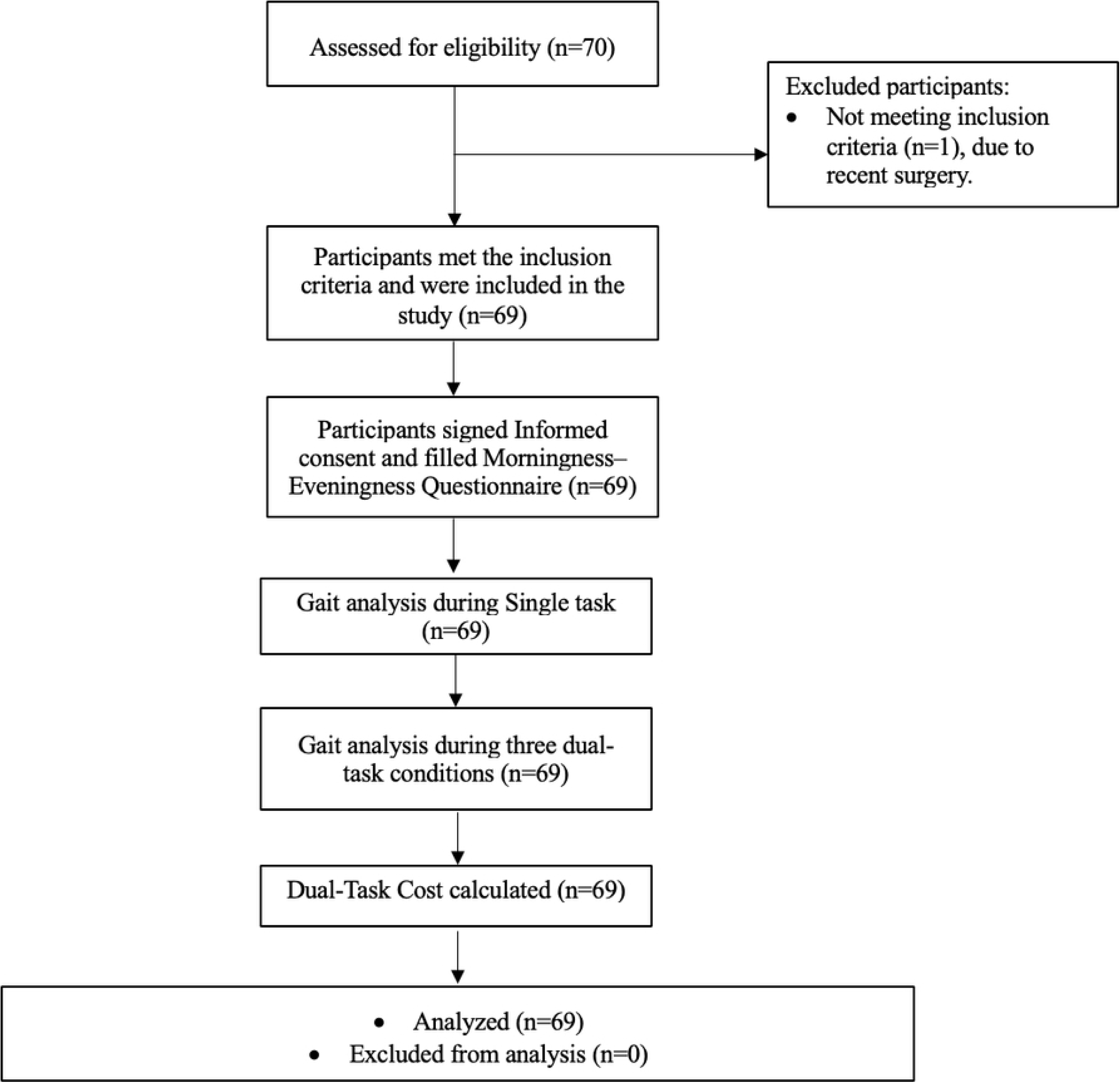
Flowchart of participants recruitment.

Inclusion criteria were: age ≥ 18 years; ability to walk independently without an assistive device; and English fluency sufficient to follow verbal instructions and complete self-report questionnaires.

Exclusion criteria were: neurological, musculoskeletal, vestibular, or balance disorder affecting gait; cardiopulmonary condition affecting mobility; use of sedating or psychoactive medications; untreated visual or sensory impairments affecting walking or task performance; and participation in organized competitive sport or structured athletic training programs. Athletes were excluded because systematic training is associated with enhanced locomotor automaticity and altered attentional strategies during walking, which could confound dual-task gait measures independently of chronotype [22,23].

### Ethical approval

The study protocol was approved by the Research Ethics Committee, University of Sharjah (Reference: REC-23-02-23-04-PG). All procedures conformed to the Declaration of Helsinki. Written informed consent was obtained from all participants prior to enrolment.

### Measures

#### Chronotype: Morningness-Eveningness Questionnaire (MEQ)

Chronotype was assessed using the Morningness–Eveningness Questionnaire (MEQ), a validated 19-item self-report instrument measuring circadian preference for sleep-wake timing and daily activity scheduling [2,24]. Scores range from 16 to 86, with higher scores indicating greater morningness [2,24]. For descriptive purposes, participants were classified into five categories: definite morning type (≥ 70), moderate morning type (59–69), intermediate type (42–58), moderate evening type (31–41), and definite evening type (≤ 30). MEQ score was treated as a continuous variable in all primary analyses to preserve statistical power and avoid arbitrary categorization. The MEQ was used under permission from the copyright holder.

#### Daytime sleepiness: Epworth Sleepiness Scale (ESS)

Daytime sleepiness was assessed using the Epworth Sleepiness Scale (ESS), an eight-item self-report questionnaire in which respondents rate their likelihood of dozing in eight common everyday situations on a 0–3 scale, yielding a total score of 0–24 [25,26]. Scores of 0–10 are within the normal range; scores above 10 indicate excessive daytime sleepiness [25,26]. The ESS was administered to screen for pathological sleepiness that could impair dual-task gait performance independently of chronotype. The ESS was administered under a licensed use agreement

#### Cognitive screening: Short Orientation-Memory-Concentration Test (SOMCT)

Global cognitive status was screened using the Short Orientation-Memory-Concentration Test (SOMCT), a brief clinician-administered instrument assessing orientation, short-term memory, and concentration [27,28]. Scores range from 0 to 28; a score of ≥ 25 is considered indicative of intact cognitive function [27,28]. The SOMCT confirmed that all participants had sufficient cognitive capacity to engage with the dual-task conditions and excluded those with undetected cognitive impairment. The SOMCT was used under permission from the copyright holder.

### Gait assessment protocol

#### Instrumentation

Spatiotemporal gait parameters were recorded using the BTS Smart Capture three-dimensional motion capture system (BTS Bioengineering, Milan, Italy), operating at a sampling rate of 100 Hz with a spatial resolution of < 1 mm [29]. The capture volume covered the full 10-metre walkway [29]. Marker trajectories were tracked by a minimum of six calibrated infrared cameras positioned to ensure continuous marker visibility throughout each trial [29].

Marker placement followed the “Helen Hayes with Medial Markers” protocol [30,31], and was performed by trained laboratory personnel prior to each session. Twenty-two Passive retroreflective markers were placed bilaterally on the following anatomical landmarks: (1) the 7th cervical vertebra; (2) bilateral acromions; (3) the 2nd sacral vertebra; (4) bilateral anterior superior iliac spines (ASIS); (5) bilateral greater trochanters; (6) bilateral medial femoral epicondyles; (7) bilateral lateral femoral epicondyles; (8) bilateral fibular heads; (9) bilateral lateral malleoli; (10) bilateral medial malleoli; (11) bilateral calcaneus; and (12) bilateral 2nd and 3rd toe junctions [23]. Anthropometric measurements—height, body weight, BMI, and bilateral leg length (anterior superior iliac spine to medial malleolus)—were recorded and used for biomechanical model scaling.

#### Single-task walking

Participants walked the 10-metre laboratory walkway at a self-selected comfortable speed. The walkway had clearly marked start and end points; participants were instructed to walk from the start to the end point and stop on arrival. No turning was performed within the measurement capture volume. Two familiarization trials were completed, and data from the third trial were used for analysis. Baseline gait velocity was included as a covariate in all regression models to account for individual differences in habitual walking speed.

The following spatiotemporal parameters were recorded: mean velocity (m/s); cadence (steps/min); stride length (m, bilateral); stride time (s, bilateral); stance time (s, bilateral); and swing time (s, bilateral).

#### Dual-task conditions

Participants performed three cognitive dual-task walking trials, each pairing the same 10-metre walking protocol with one of the following cognitive tasks:

##### Backward word spelling (5LWB)

Participants spelled a five-letter word backwards aloud while walking. Words were presented verbally by the assessor immediately before each trial. This task engages phonological processing, orthographic retrieval, response inhibition of habitual forward spelling, and verbal working memory [22,23].

##### Serial subtraction by seven (SS7)

Participants counted backwards from 100 by subtracting seven from each successive number aloud while walking. This task engages arithmetic processing, working memory updating, and sustained attention [22,23].

##### Reverse month recitation (RMR)

Participants recited the months of the year in reverse order from a randomly assigned starting month aloud while walking. This task requires sequential processing, inhibition of habitual forward recitation, and working memory [22,23,32].

These three tasks were selected because they represent distinct cognitive processing domains—phonological, arithmetic, and sequential—enabling comparison of domain-specific dual-task interference effects. All three are established cognitive probes used in prior dual-task gait research and can be administered without specialized equipment [22,23].

Each task was explained and demonstrated by the assessor before the trial. Participants were instructed to begin the cognitive task simultaneously with their first step and to continue both tasks until reaching the end of the walkway. No explicit prioritization instruction was given; participants were asked to maintain their natural walking pace while performing the cognitive task, consistent with standard dual-task gait protocols [33,34]. Verbal responses were monitored by the assessor to confirm continuous task engagement. Cognitive task accuracy and response rates were not formally recorded; this is acknowledged as a limitation of the present study.

The order of the three dual-task conditions was randomized across participants using a computer-generated counterbalanced sequence (block size 6) to control for order and fatigue effects. Rest periods of at least three minutes were provided between trials to minimize fatigue.

#### Dual-task cost (DTC)

Dual-task cost was calculated for each gait parameter using:

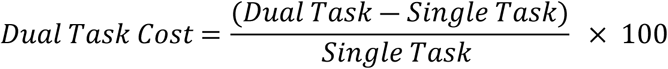

A negative DTC for velocity, cadence, and stride length indicates performance deterioration; a positive DTC for stride time, stance time, and swing time indicates slower, more cautious gait. This sign convention was applied consistently across all parameters [12,31,35,36].

### Statistical analysis

All analyses were conducted using IBM SPSS Statistics Version 25 (IBM Corp., Armonk, NY, USA). Statistical significance was set at p < 0.05. Data are reported as mean ± standard deviation unless otherwise stated.

Normality was assessed using the Shapiro-Wilk test and Q-Q plot inspection. Non-normal DTC distributions were analyzed using non-parametric tests. Continuous predictors (MEQ score, age, BMI, baseline gait velocity) were mean-centered prior to regression to improve interpretability. Multicollinearity was assessed using variance inflation factors (VIF); VIF < 2 indicated no collinearity concern.

Differences in DTC across the three dual-task conditions (5LWB, SS7, RMR) were evaluated using the Friedman test for each gait parameter. Statistically significant Friedman results were followed by pairwise Wilcoxon signed-rank post-hoc tests across the three condition pairs (5LWB vs. SS7, 5LWB vs. RMR, SS7 vs. RMR), with Bonferroni correction applied to control the family-wise error rate (adjusted α = 0.017).

To evaluate the independent association between chronotype and dual-task gait performance, separate multiple linear regression models were constructed for each gait parameter and each task condition. The dependent variable was DTC (%) for each gait parameter; the primary predictor was MEQ total score (continuous); covariates were age, sex, BMI, and the corresponding single-task baseline gait value. Mean velocity DTC is the primary outcome; all remaining gait parameters are secondary. Standardized regression coefficients (β), model R², and p-values are reported.

Spearman rank correlation coefficients (ρ) were calculated to examine bivariate associations between MEQ score and DTC across all gait parameters and conditions. These analyses were pre-specified as exploratory. Given the number of comparisons, secondary parameter-level findings should be interpreted with appropriate caution.

### Data availability

All data supporting the findings of this study are publicly available on Zenodo: https://doi.org/10.5281/zenodo.17878983

## Results

### Participant characteristics

A total of 69 participants (37 females, 32 males) with a mean age of 21.28 ± 2.83 years (range 18–35) were included. The mean MEQ score was 46.16 ± 8.62, corresponding to a predominantly intermediate chronotype distribution: 45 participants were intermediate types (65.2%), 18 were moderate evening types (26.1%), and five were morning types (7.2%; three moderate morning, two definite morning). Mean baseline gait velocity was 1.18 ± 0.13 m/s. Daytime sleepiness (ESS mean: 8.74 ± 4.05) and cognitive screening (SOMCT mean: 26.81 ± 1.13 out of 28) were within normal ranges for all participants. Baseline demographic characteristics and gait parameters are summarized in **Table 1**.

**Table 1.**
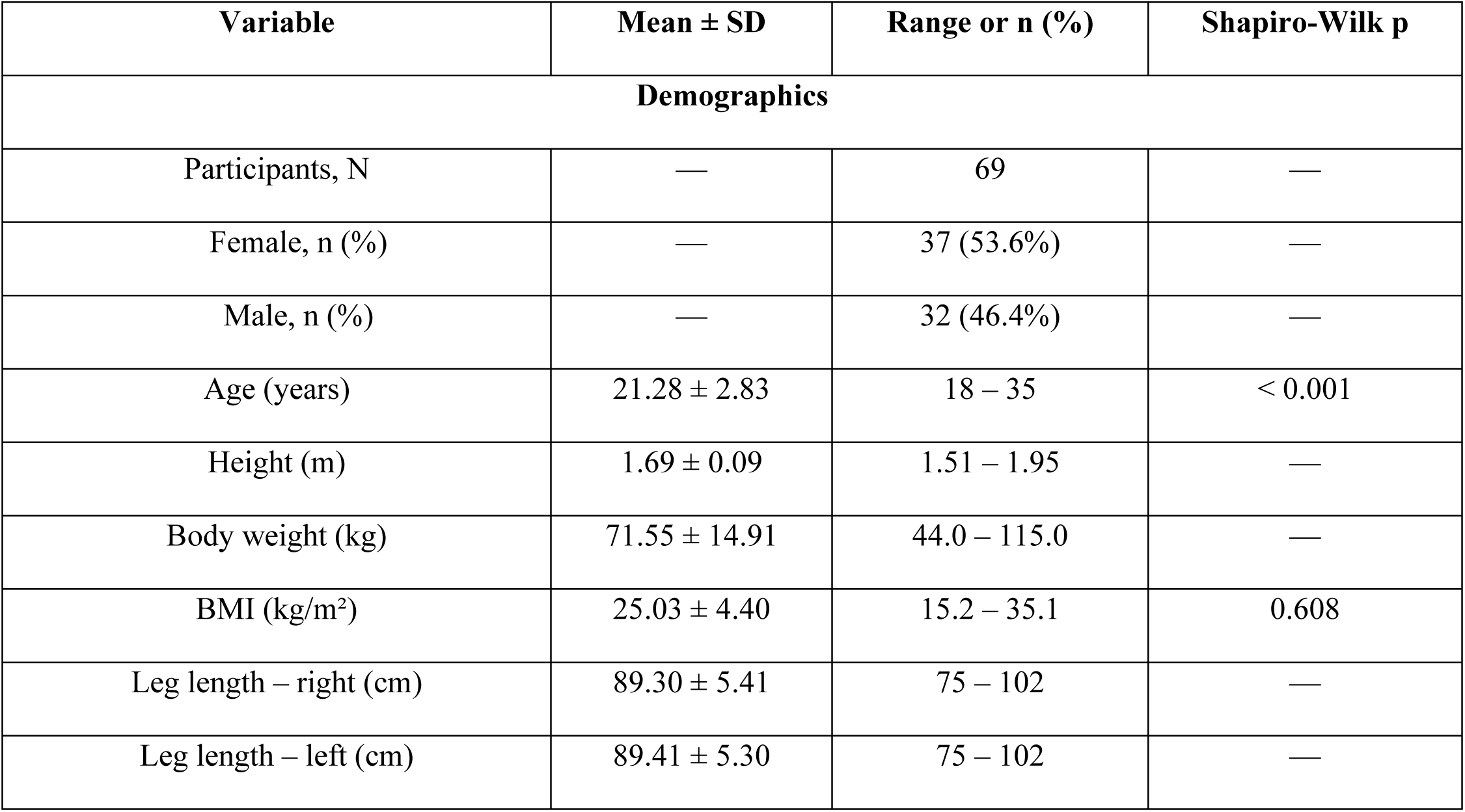
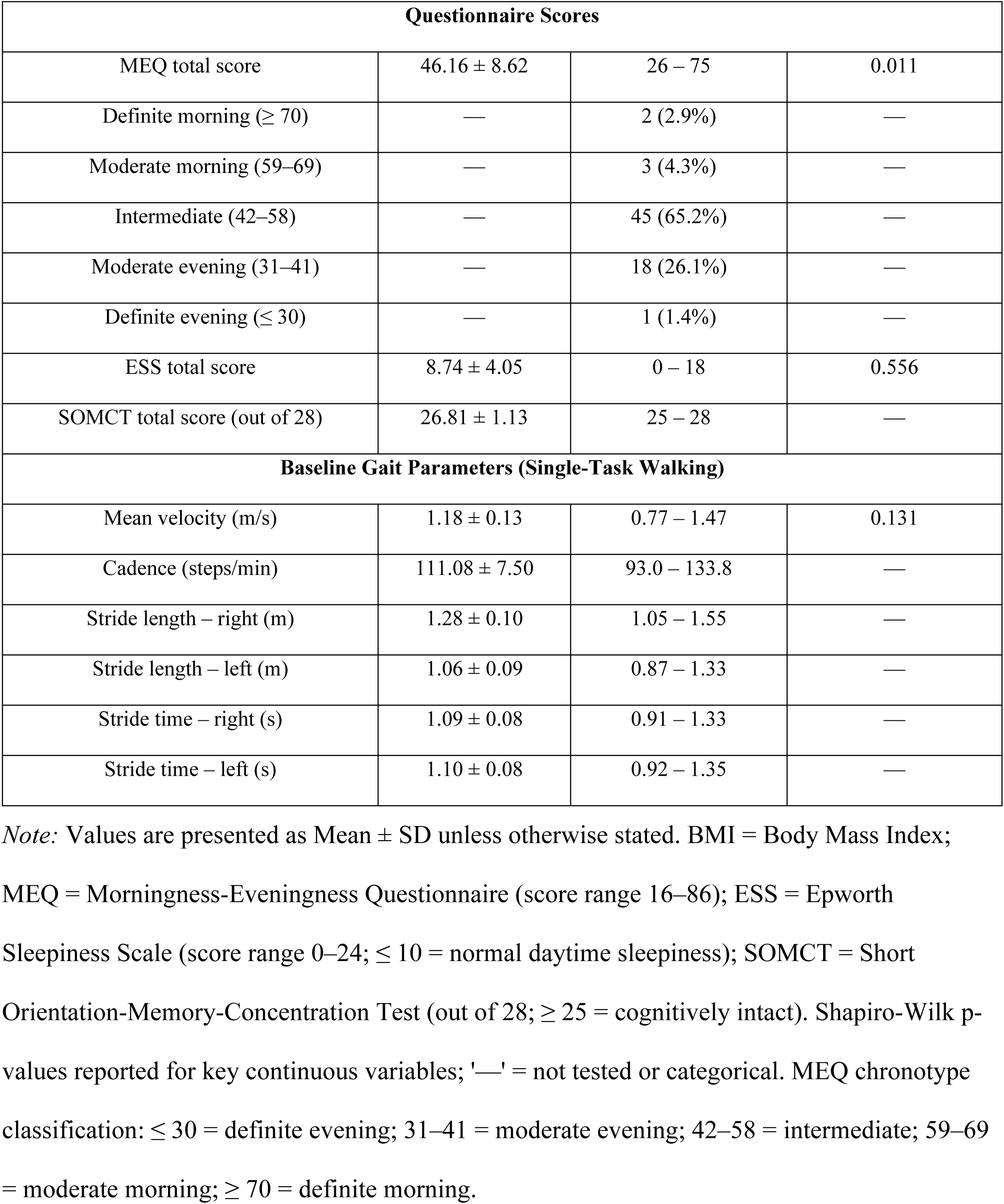
Demographic Characteristics, Questionnaire Scores, and Baseline Gait Parameters.

| Variable | Mean ± SD | Range or n (%) | Shapiro-Wilk p |
| --- | --- | --- | --- |
| **Demographics** |  |  |  |
| Participants, N | — | 69 | — |
| Female, n (%) | — | 37 (53.6%) | — |
| Male, n (%) | — | 32 (46.4%) | — |
| Age (years) | $21.28 \pm 2.83$ | 18 – 35 | $< 0.001$ |
| Height (m) | $1.69 \pm 0.09$ | 1.51 – 1.95 | — |
| Body weight (kg) | $71.55 \pm 14.91$ | 44.0 – 115.0 | — |
| BMI (kg/m<sup>2</sup>) | $25.03 \pm 4.40$ | 15.2 – 35.1 | 0.608 |
| Leg length – right (cm) | $89.30 \pm 5.41$ | 75 – 102 | — |
| Leg length – left (cm) | $89.41 \pm 5.30$ | 75 – 102 | — |

| Questionnaire Scores |  |  |  |
| --- | --- | --- | --- |
| MEQ total score | 46.16 ± 8.62 | 26 – 75 | 0.011 |
| Definite morning (≥ 70) | — | 2 (2.9%) | — |
| Moderate morning (59–69) | — | 3 (4.3%) | — |
| Intermediate (42–58) | — | 45 (65.2%) | — |
| Moderate evening (31–41) | — | 18 (26.1%) | — |
| Definite evening (≤ 30) | — | 1 (1.4%) | — |
| ESS total score | 8.74 ± 4.05 | 0 – 18 | 0.556 |
| SOMCT total score (out of 28) | 26.81 ± 1.13 | 25 – 28 | — |
| Baseline Gait Parameters (Single-Task Walking) |  |  |  |
| Mean velocity (m/s) | 1.18 ± 0.13 | 0.77 – 1.47 | 0.131 |
| Cadence (steps/min) | 111.08 ± 7.50 | 93.0 – 133.8 | — |
| Stride length – right (m) | 1.28 ± 0.10 | 1.05 – 1.55 | — |
| Stride length – left (m) | 1.06 ± 0.09 | 0.87 – 1.33 | — |
| Stride time – right (s) | 1.09 ± 0.08 | 0.91 – 1.33 | — |
| Stride time – left (s) | 1.10 ± 0.08 | 0.92 – 1.35 | — |

*Note:* Values are presented as Mean ± SD unless otherwise stated. BMI = Body Mass Index; MEQ = Morningness-Eveningness Questionnaire (score range 16–86); ESS = Epworth Sleepiness Scale (score range 0–24; ≤ 10 = normal daytime sleepiness); SOMCT = Short Orientation-Memory-Concentration Test (out of 28; ≥ 25 = cognitively intact). Shapiro-Wilk p-values reported for key continuous variables; '—' = not tested or categorical. MEQ chronotype classification: ≤ 30 = definite evening; 31–41 = moderate evening; 42–58 = intermediate; 59–69 = moderate morning; ≥ 70 = definite morning.

### Dual-task cost across conditions

Shapiro-Wilk testing indicated non-normal DTC distributions for 7 of 8 gait parameters under dual-task conditions, supporting use of non-parametric analyses. The Friedman test revealed significant differences in DTC across the three cognitive conditions for all eight spatiotemporal gait parameters (all p < 0.05; Table 2).

**Table 2.**
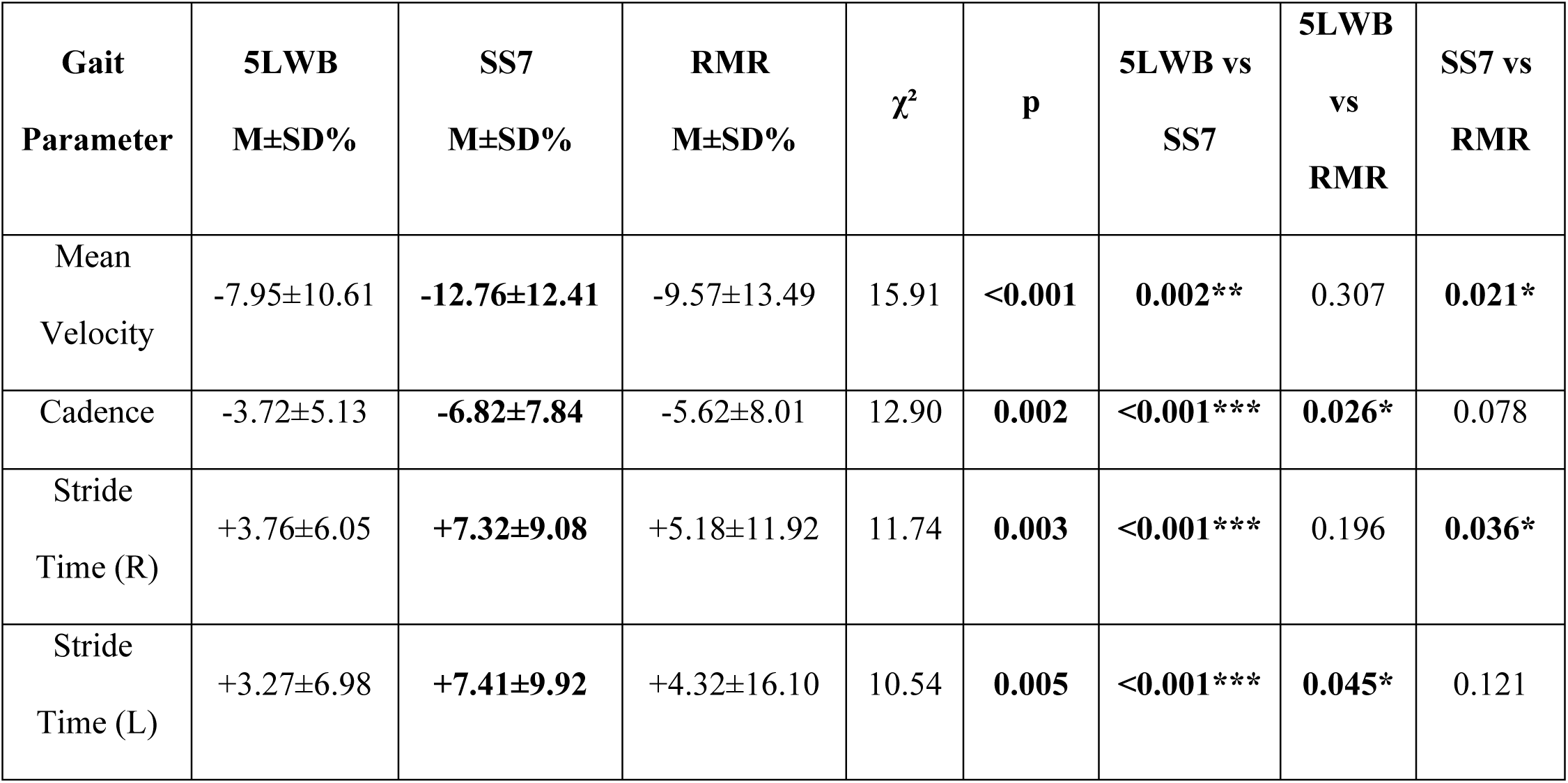
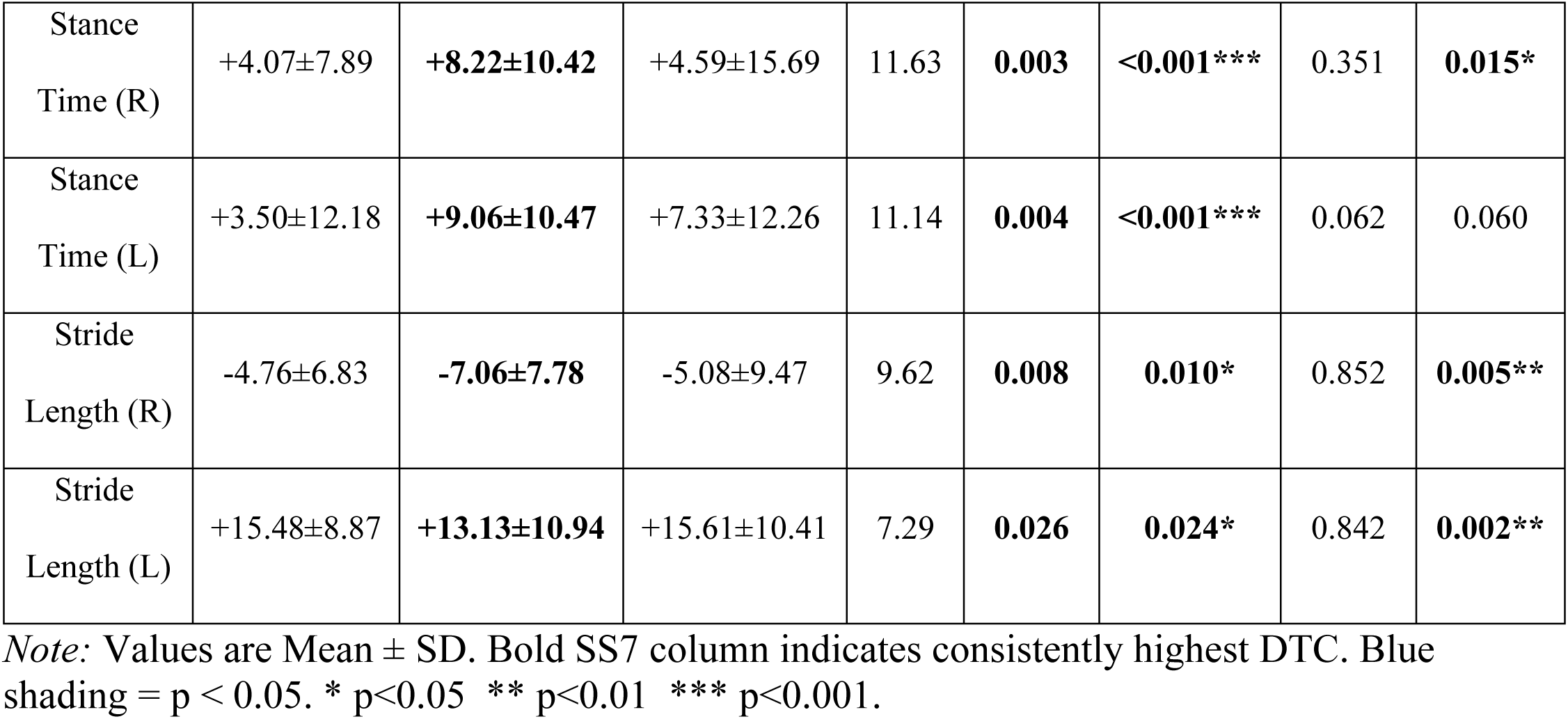
Friedman test and Wilcoxon signed-rank post-hoc comparisons of DTC (%) across cognitive conditions.

| Gait Parameter | 5LWB<br>M±SD% | SS7<br>M±SD% | RMR<br>M±SD% | $\chi^2$ | p | 5LWB vs<br>SS7 | 5LWB<br>vs<br>RMR | SS7 vs<br>RMR |
| --- | --- | --- | --- | --- | --- | --- | --- | --- |
| Mean Velocity | -7.95±10.61 | **-12.76±12.41** | -9.57±13.49 | 15.91 | **<0.001** | **0.002\*\*** | 0.307 | **0.021\*** |
| Cadence | -3.72±5.13 | **-6.82±7.84** | -5.62±8.01 | 12.90 | **0.002** | **<0.001\*\*\*** | **0.026\*** | 0.078 |
| Stride Time (R) | +3.76±6.05 | **+7.32±9.08** | +5.18±11.92 | 11.74 | **0.003** | **<0.001\*\*\*** | 0.196 | **0.036\*** |
| Stride Time (L) | +3.27±6.98 | **+7.41±9.92** | +4.32±16.10 | 10.54 | **0.005** | **<0.001\*\*\*** | **0.045\*** | 0.121 |
| Stance<br>Time (R) | +4.07±7.89 | **+8.22±10.42** | +4.59±15.69 | 11.63 | **0.003** | **<0.001\*\*\*** | 0.351 | **0.015\*** |
| Stance<br>Time (L) | +3.50±12.18 | **+9.06±10.47** | +7.33±12.26 | 11.14 | **0.004** | **<0.001\*\*\*** | 0.062 | 0.060 |
| Stride<br>Length (R) | -4.76±6.83 | **-7.06±7.78** | -5.08±9.47 | 9.62 | **0.008** | **0.010\*** | 0.852 | **0.005\*\*** |
| Stride<br>Length (L) | +15.48±8.87 | **+13.13±10.94** | +15.61±10.41 | 7.29 | **0.026** | **0.024\*** | 0.842 | **0.002\*\*** |

*Note:* Values are Mean ± SD. Bold SS7 column indicates consistently highest DTC. Blue shading = $p < 0.05$ . \* $p < 0.05$ \*\* $p < 0.01$ \*\*\* $p < 0.001$ .

For mean velocity DTC, a significant overall difference was found across conditions (Friedman χ² = 15.91, p < 0.001). The greatest mean velocity DTC was observed during the SS7 condition (−12.76 ± 12.41%), followed by RMR (−9.57 ± 13.49%) and 5LWB (−7.95 ± 10.61%). Post-hoc pairwise comparisons (Bonferroni-corrected Wilcoxon signed-rank) showed that SS7 produced significantly greater DTC than both 5LWB (p = 0.002) and RMR (p = 0.021); the 5LWB vs. RMR difference was not significant (p = 0.307). This pattern was replicated across all eight gait parameters, as shown in **Table 2**. Boxplots with individual data points are presented in **Fig 2**.

**Fig 2.**
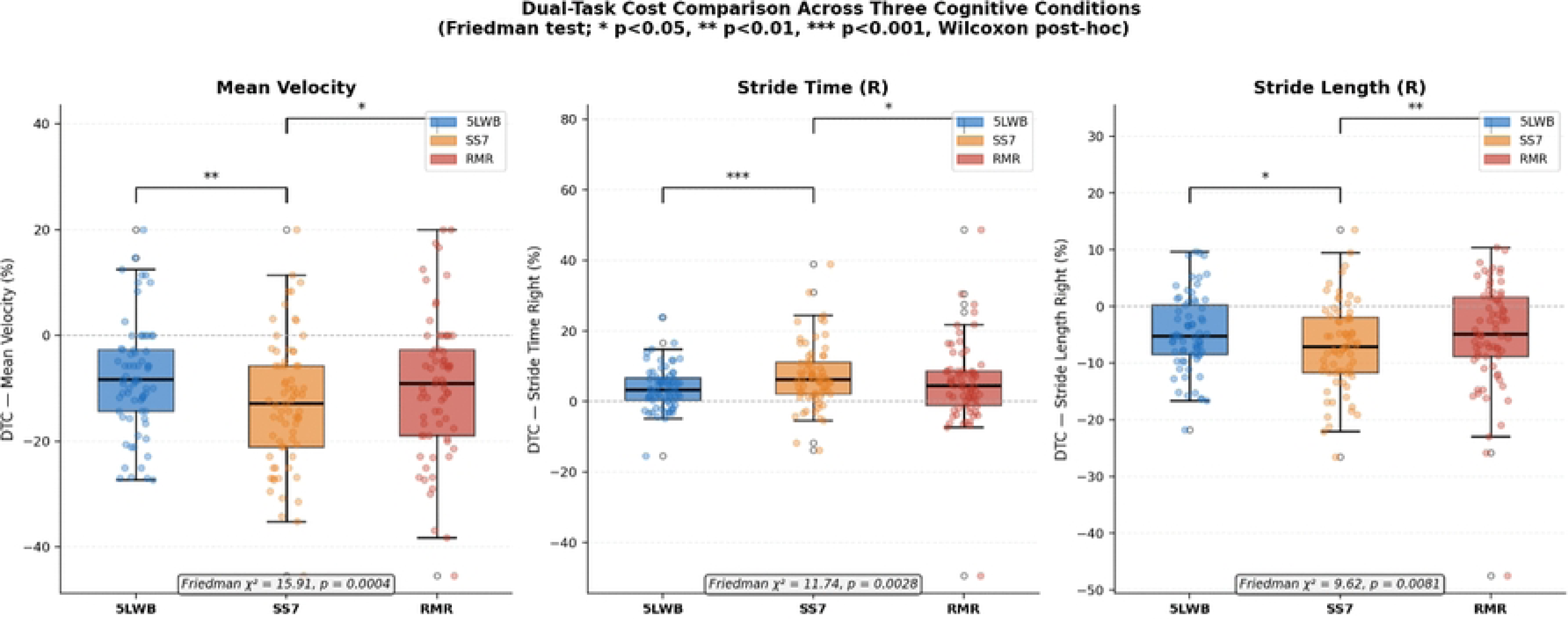
Boxplots comparing DTC across 5LWB, SS7, and RMR conditions for mean velocity, stride time (right), and stride length (right). Individual data points overlaid. Significance brackets: Wilcoxon signed-rank (* p<0.05, ** p<0.01, *** p<0.001). Friedman statistics shown per panel.

### Association between chronotype and dual-task cost

Multiple linear regression analyses were conducted to examine the independent association between MEQ score and mean velocity DTC across the three conditions, adjusting for age, sex, BMI, and baseline gait velocity (N = 69). All variance inflation factors were below 2 (MEQ = 1.13, Age = 1.29, Sex = 1.30, BMI = 1.10, baseline velocity = 1.06), confirming the absence of multicollinearity.

In the 5LWB condition, MEQ score was a significant independent predictor of mean velocity DTC (β = −0.51, p < 0.001, R² = 0.269), indicating that higher morningness was independently associated with better preservation of gait speed under phonological cognitive load. In the RMR condition, MEQ was also a significant predictor (β = −0.55, p = 0.004, R² = 0.222). In the SS7 condition, the association did not reach statistical significance (β = −0.33, p = 0.071, R² = 0.118), though the direction was consistent. Regression results across all gait parameters and conditions are presented in **Table 3**. Scatter plots with regression lines and 95% confidence intervals are shown in **Fig 3**.

**Fig 3.**
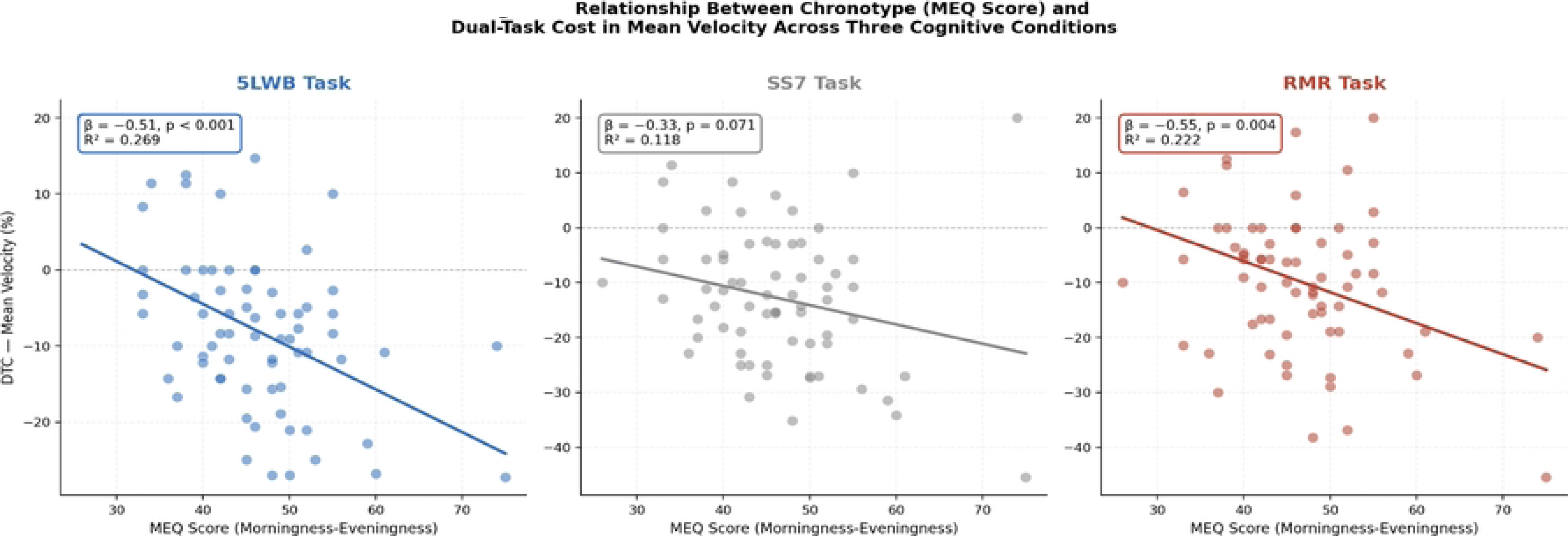
Scatter plots of MEQ score vs dual-task cost in mean velocity for 5LWB (left), SS7 (middle), and RMR (right). Regression lines with 95% confidence intervals shown. β and p-values are from multiple regression controlling for age, sex, BMI, and baseline velocity.

**Table 3.** Multiple linear regression between MEQ score and mean velocity DTC across the three conditions, adjusting for age, sex, BMI, and baseline gait velocity.

| Gait Parameter | 5LWB $\beta$ | 5LWB p | 5LWB $R^2$ | SS7 $\beta$ | SS7 p | SS7 $R^2$ | RMR $\beta$ | RMR p | RMR $R^2$ |
| --- | --- | --- | --- | --- | --- | --- | --- | --- | --- |
| Mean Velocity | **-0.512** | **<0.001\*\*\*** | 0.269 | -0.333 | 0.071 | 0.118 | **-0.553** | **0.004\*\*** | 0.222 |
| Cadence | **-0.162** | **0.038\*** | 0.078 | -0.077 | 0.523 | 0.022 | **-0.281** | **0.019\*** | 0.127 |
| Stride Time (R) | **+0.227** | **0.011\*** | 0.154 | +0.237 | 0.086 | 0.069 | **+0.610** | **<0.001\*\*\*** | 0.250 |
| Stride Time (L) | **+0.327** | **0.002\*\*** | 0.171 | +0.122 | 0.428 | 0.027 | +0.422 | 0.077 | 0.124 |
| Stance Time (R) | **+0.266** | **0.023\*** | 0.140 | +0.187 | 0.240 | 0.056 | -0.072 | 0.759 | 0.084 |
| Stance Time (L) | -0.141 | 0.456 | 0.020 | +0.285 | 0.076 | 0.064 | **+0.606** | **<0.001\*\*\*** | 0.223 |
| Stride Length (R) | **-0.218** | **0.030\*** | 0.150 | -0.220 | 0.054 | 0.144 | -0.162 | 0.217 | 0.222 |
| Stride<br>Length (L) | **-0.354** | **0.005\*\*** | 0.212 | **-0.378** | **0.018\*** | 0.156 | **-0.341** | **0.024\*** | 0.167 |

*Note:* MEQ $\beta$ coefficients, p-values, and $R^2$ for DTC across all gait parameters and tasks.
Covariates: age, sex, BMI, baseline walking velocity (N = 69). Blue shading = $p < 0.05$ . \*
$p < 0.05$ \*\* $p < 0.01$ \*\*\* $p < 0.001$ .

Baseline gait velocity was the only covariate reaching statistical significance, and only in the RMR model (p = 0.040). Age, sex, and BMI were non-significant across all three models.

### Exploratory Spearman correlations

Spearman correlations between MEQ score and mean velocity DTC were significant across all three conditions: 5LWB (ρ = −0.385, p = 0.001), SS7 (ρ = −0.257, p = 0.033), and RMR (ρ = −0.281, p = 0.019). Across all eight gait parameters, the number of significant MEQ–DTC associations was task-dependent: 7 of 8 parameters in 5LWB, 4 of 8 in RMR, and 3 of 8 in SS7. Non-significant associations were most prevalent in the SS7 condition, particularly for temporal gait parameters (stride time, stance time, swing time). A heatmap of regression β coefficients across all gait parameters and conditions is presented in **Fig 4**.

**Fig 4.**
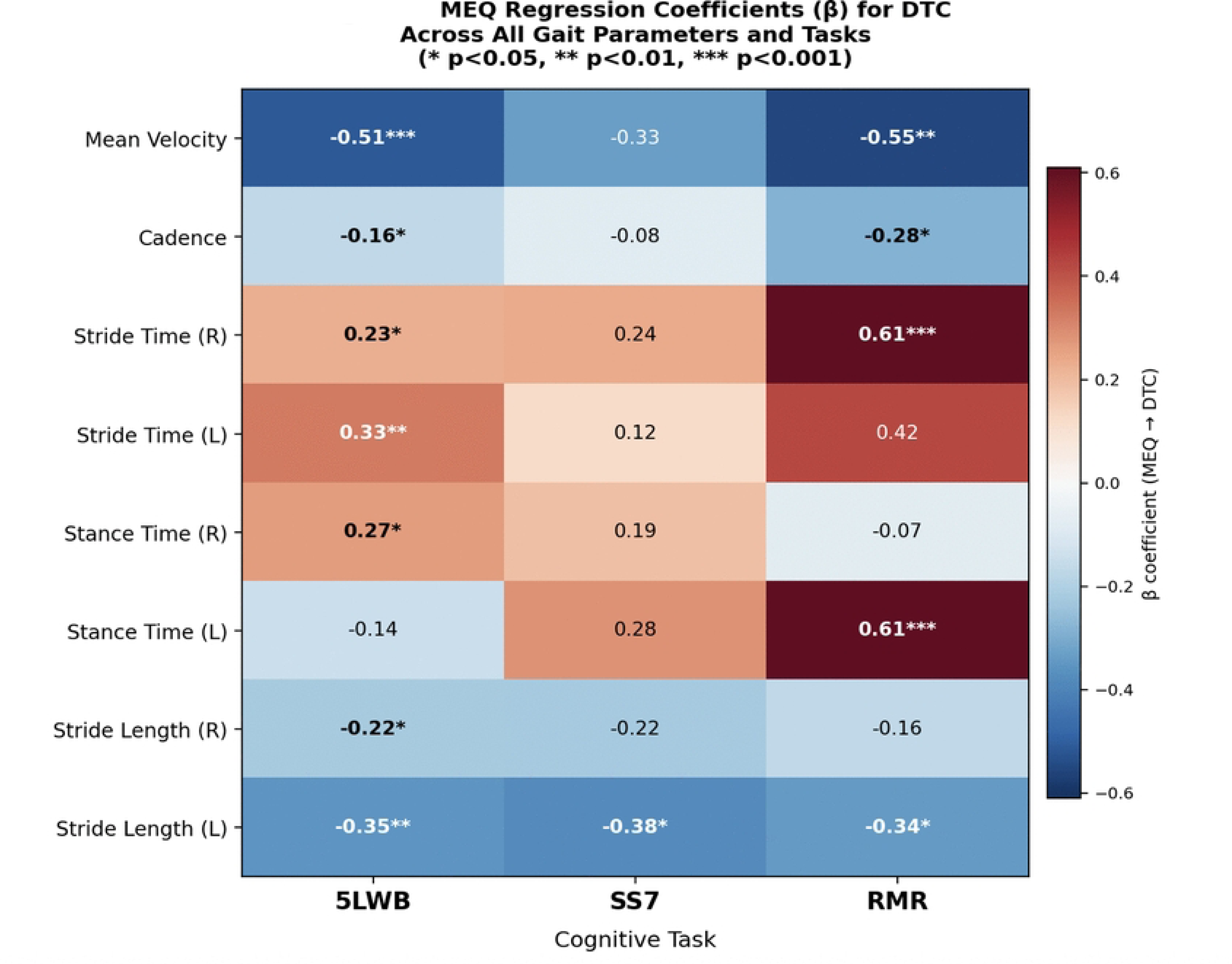
Heatmap of MEQ regression β coefficients across all gait parameters and tasks. Blue = negative association (morningness protective); Red = positive association (greater DTC with higher morningness, reflecting relative time-variable adaptation). * p<0.05 ** p<0.01 *** p<0.001.

## Discussion

This study examined the association between chronotype and dual-task gait performance in young adults across three distinct cognitive conditions. The primary finding is that chronotype is independently associated with dual-task gait cost in a task-dependent manner, with consistent effects observed in the phonological (5LWB) and sequential (RMR) conditions, but not in the arithmetic (SS7) condition.

### Chronotype and dual-task gait: task-dependent effects

Multiple regression analyses, adjusting for age, sex, BMI, and baseline gait velocity, demonstrated that MEQ score was a significant independent predictor of dual-task cost in mean velocity during the 5LWB (β = −0.51, p < 0.001, R² = 0.269) and RMR (β = −0.55, p = 0.004, R² = 0.222) conditions. In both cases, higher MEQ scores—indicating greater morningness—were associated with better preservation of gait speed under cognitive load, suggesting more efficient cognitive–motor integration under these specific task demands. All VIFs were below 2, confirming the absence of multicollinearity and the stability of these estimates.

No significant independent association was observed in the SS7 condition (β = −0.33, p = 0.071), though the direction was consistent. This divergence indicates that chronotype does not exert a uniform effect across all forms of cognitive interference but rather depends on the nature of the concurrent cognitive task. Exploratory Spearman correlations corroborated this gradient: significant MEQ–DTC associations were detected in 7 of 8 gait parameters during 5LWB, 4 of 8 during RMR, and only 3 of 8 during SS7.

A plausible explanation lies in the differing neural demands of the tasks. The 5LWB task engages phonological processing, response inhibition, and executive control, while the RMR task requires sequential processing and working memory updating. Both domains are closely linked to prefrontal cortical networks—including the dorsolateral prefrontal cortex (DLPFC) and anterior cingulate cortex (ACC)—which are known to exhibit circadian modulation and to overlap substantially with the neural circuits supporting gait control under attentional challenge [37,38]. In contrast, the SS7 task relies more heavily on arithmetic processing, which may depend on partially distinct parietal circuits—including the intraparietal sulcus and angular gyrus [39,40]—less sensitive to circadian variation in arousal. This may explain the weaker chronotype effects in the SS7 condition. These neural interpretations remain speculative and should be interpreted with caution, as no neurophysiological measures were collected in the present study.

### Dual-task interference across cognitive conditions

Independent of chronotype, the Friedman test confirmed that task identity significantly modulated gait interference across all eight spatiotemporal parameters (all p < 0.05). The SS7 condition produced the greatest overall dual-task cost, with mean velocity DTC of −12.76 ± 12.41%, significantly exceeding both 5LWB and RMR.

This finding contrasts with the pattern observed for chronotype effects, where SS7 showed the fewest significant associations. Together, these results indicate that overall task difficulty, as reflected by absolute gait disruption, and chronotype sensitivity, as reflected by individual variability in dual-task cost, represent distinct constructs and may not be driven by the same underlying mechanisms. This interpretation remains inferential and should not be considered evidence of distinct neural pathways without direct neurophysiological validation.

That SS7 failed to show chronotype sensitivity despite producing the largest absolute DTC may reflect a ceiling effect on gait disruption, or alternatively that arithmetic-driven interference is less amenable to circadian arousal modulation. However, this interpretation remains speculative and requires confirmation in studies incorporating neurophysiological measures or within-subject circadian designs [3].

In the absence of concurrent cognitive performance data, it cannot be determined whether participants prioritized gait or cognitive task performance across conditions. The absence of cognitive task performance measures (accuracy and response rates) limits interpretation of dual-task trade-offs. It is therefore not possible to determine whether participants prioritized gait or cognitive performance, and the observed dual-task cost may reflect task prioritization strategies rather than pure cognitive–motor interference [41].

### Interpretation of temporal versus spatial gait changes

An important pattern in the regression analyses is the differential direction of MEQ associations across gait domains. Higher MEQ scores were associated with reduced DTC in velocity and stride length, indicating better spatial gait preservation, while simultaneously being associated with increased DTC in temporal parameters such as stride time and stance time in several conditions.

This apparent discrepancy likely reflects an adaptive motor control strategy rather than contradictory effects. Increased temporal DTC—longer stride or stance durations under dual-task conditions—can indicate a more cautious gait pattern that allows individuals to maintain stability while preserving overall spatial performance. In this context, morning-type individuals may adopt compensatory temporal adjustments that enable better maintenance of gait velocity under cognitive load. The observed pattern may therefore reflect a shift toward stability-oriented motor control rather than performance deterioration and is consistent with evidence that dual-task gait adaptations in healthy young adults are not uniformly maladaptive [42,43].

### Role of baseline gait and covariates

Baseline gait velocity emerged as a significant covariate only in the RMR model (p = 0.040), indicating that initial motor capacity may specifically influence dual-task performance under sequential cognitive demands. Age, sex, and BMI were non-significant in all three models, suggesting that the observed chronotype effects are not attributable to basic demographic or anthropometric differences in this sample.

### Chronotype, time-of-day, and cognitive arousal

The observed association between morningness and lower DTC is consistent with the synchrony hypothesis, which predicts that cognitive performance is optimized when task timing aligns with an individual’s circadian peak [6,7,44]. Morning types generally reach peak alertness and executive function in the late morning to early afternoon [7,8,44], which corresponds to the testing window used in this study (12:00–16:00). Evening types typically experience delayed arousal trajectories during this period, which could reduce the prefrontal resources available for simultaneous cognitive and gait management [45,46].

However, the fixed testing window does not allow causal inference regarding chronotype effects independent of circadian phase. As all participants were assessed within the same time period, the observed associations may reflect a combined influence of trait chronotype and time-of-day–specific physiological state, rather than chronotype alone [6,7,44].

This limitation represents the primary constraint on interpretation of the present findings and precludes definitive conclusions regarding chronotype-specific mechanisms, particularly given that higher-order cognitive functions mediated by prefrontal networks are known to be sensitive to circadian modulation [8].

Additional individual-difference factors that were not captured by the MEQ or ESS—including sleep quality, habitual sleep duration, trait anxiety, and motivation—may also contribute to the observed pattern and cannot be excluded as explanations with the present data [42,47].

### Methodological considerations and limitations

Several limitations should be considered when interpreting these findings. First, and most critically, all assessments were conducted at a fixed time of day without alignment to participants’ individual circadian preferences. This design confounds chronotype with time-of-day effects and is the primary constraint on causal interpretation, as discussed above.

Second, cognitive task performance metrics—accuracy and response rates—were not recorded during the dual-task walking trials. This limits the ability to assess cognitive–motor performance trade-offs, and means that observed gait changes cannot be unambiguously attributed to dual-task interference rather than strategic cognitive task prioritization.

Third, the sample was predominantly composed of intermediate chronotypes (65.2%), with very few morning-type participants (n = 5 combined). This distribution, consistent with the documented epidemiology of chronotype in young adults [4,5], reduces statistical power to detect associations at the extremes of the chronotype spectrum and means results most accurately characterize intermediate-to-moderate-evening types.

Fourth, given the number of statistical tests performed across multiple gait parameters and conditions, secondary findings—particularly individual parameter-level correlations—should be interpreted with appropriate caution.

Finally, chronotype was measured exclusively via the MEQ, a validated but subjective self-report instrument. Objective circadian phase estimation using actigraphy or salivary dim-light melatonin onset (DLMO) profiling would strengthen biological validity in future work [3].

### Implications and future directions

Despite these limitations, the present findings provide evidence that chronotype is associated with dual-task gait performance in a cognitively specific manner, rather than as a general effect across all task types. This contributes to understanding of individual variability in cognitive–motor interference and suggests that circadian factors may selectively influence executive and sequential processing during locomotion.

These findings do not yet support recommendations for chronotype-based scheduling in clinical rehabilitation settings, as the sample comprised healthy young adults and the design does not establish causality. Rather, they identify chronotype as a variable warranting inclusion as a covariate in cognitive-motor research, and support the following priorities for future investigation: (1) within-participant multi-time-point designs to isolate circadian synchrony effects from trait chronotype; (2) concurrent recording of cognitive task performance to enable trade-off analysis; (3) recruitment of balanced chronotype groups including larger morning-type samples; and (4) neurophysiological approaches to examine the proposed prefrontal mechanisms.

## Conclusions

Chronotype is associated with dual-task gait cost in a task-domain-specific manner in healthy young adults. Morningness was associated with better gait speed preservation under phonological and sequential cognitive loads, consistent with the circadian synchrony hypothesis. In contrast, no significant chronotype effect was observed in arithmetic conditions. These findings suggest a potential association between chronotype and dual-task gait performance; however, causal relationships cannot be established due to the fixed testing window and cross-sectional design. Future research using within-subject time-of-day–controlled designs, concurrent cognitive performance recording, and objective circadian phase markers is needed to confirm these associations and clarify the underlying mechanisms.

## Author contributions

Conceptualisation: MK, JD, AJAS. Methodology: MK, JD, AJAS. Data curation and formal analysis: MK, JD. Investigation: JD, MK. Writing – original draft: MK, JD. Writing – review and editing: AJAS. Supervision: MK, AJAS. All authors read and approved the final manuscript.

